# The effect of magnesium on calcium binding to cardiac troponin C related hypertrophic cardiomyopathy mutants

**DOI:** 10.1101/2021.05.12.443789

**Authors:** Kaveh Rayani, Eric Hantz, Omid Haji-Ghassemi, Alison Yueh Li, Anne Marie Spuches, Filip Van Petegem, R John Solaro, Steffen Lindert, Glen F Tibbits

## Abstract

Cardiac troponin C (cTnC) is the calcium (Ca^2+^) sensing component of the troponin complex. Binding of Ca^2+^ to cTnC triggers a cascade of myofilament conformational changes that culminate in force production. Mutations in cTnC linked to hypertrophic myocardial myopathy (HCM) induce a a greater degree and duration of Ca^2+^ binding, which may underly the hypertrophic phenotype. Recent evidence from our laboratories demonstrated novel modifications of cTnC Ca^2+^ binding by cellular magnesium (Mg^2+^) that we hypothesize may be of significance in promoting HCM.

Regulation of contraction has long been thought to occur exclusively through Ca^2+^ binding to site II of cTnC. However, abundant cellular Mg^2+^ is a potential competitor for binding to the same sites; work by several groups also suggests this is possible. We have used isothermal titration calorimetry (ITC) to explore the thermodynamic properties associated with the interaction between Ca^2+^/Mg^2+^ and site II of cTnC; these experiments demonstrated that physiological concentrations of Mg^2+^ may compete with Ca^2+^ to bind site II of cTnC.

In experiments reported here, we studied a series of mutations in cTnC thought to be causal in HCM. Three mutants (A8V, L29Q, and A31S) slightly elevated the affinity for both Ca^2+^ and Mg^2+^, whereas other mutants (L48Q, Q50R, and C84Y), that are closer to the C-terminal domain and surrounding the EF hand binding motif of site II had a more significant effect on affinity and the thermodynamics of the binding interaction.

To the best of our knowledge, this work is the first to explore the role of Mg^2+^ in modifying the Ca^2+^ affinity ofcTnC mutations linked to HCM. Our results indicate a physiologically significant role for cellular Mg^2+^ at baseline conditions and when elevated on the control of the dynamics of contraction by modifications in the Ca^2+^ binding properties of cTnC.

## Introduction

The studies reported here compare Mg^2+^-induced modifications in the Ca^2+^ binding properties of the regulatory site of cardiac Troponin C (cTnC) with mutations linked to cardiomyopathies. cTnC is a globular protein with 4 EF hands domains. High affinity structural sites III and IV in the C-terminal domain bind Ca^2+^ (K_d_ ~ 0.1 μM) and Mg^2+^ (K_d_ ~100 μM) to tether cTnC to the other components of the cTn complex (1–3). The N-terminal domain contains site I which is dysfunctional in cardiac muscle, and site II which plays a regulatory role. Low affinity (K_A_ ~10^5^ M^-1^) site II binds Ca^2+^ at elevated concentrations during systolic (~1 μM) and is unbound during diastole (~100 nM) (4). cTnC is a dumbbell-like protein composed of nine α-helices helices. Helices N, A, B, C, and D in the N-domain are linked through the flexible D-E linker to the C-domain which contains helices E, F, G, and H. Binding of Ca^2+^ to site II acts as a conformational switch in cTnC, with the helices N, A and D moving away from helices B and C, exposing a hydrophobic cleft (**Figure 1**). This region is then bound by the cTnI switch peptide (TnI_sw_), causing further perturbation within the cTn complex and the rest of the TF to expose actin binding sites allowing for contact with myosin heads, ultimately resulting in force production (5,6).

**Figure 1:**
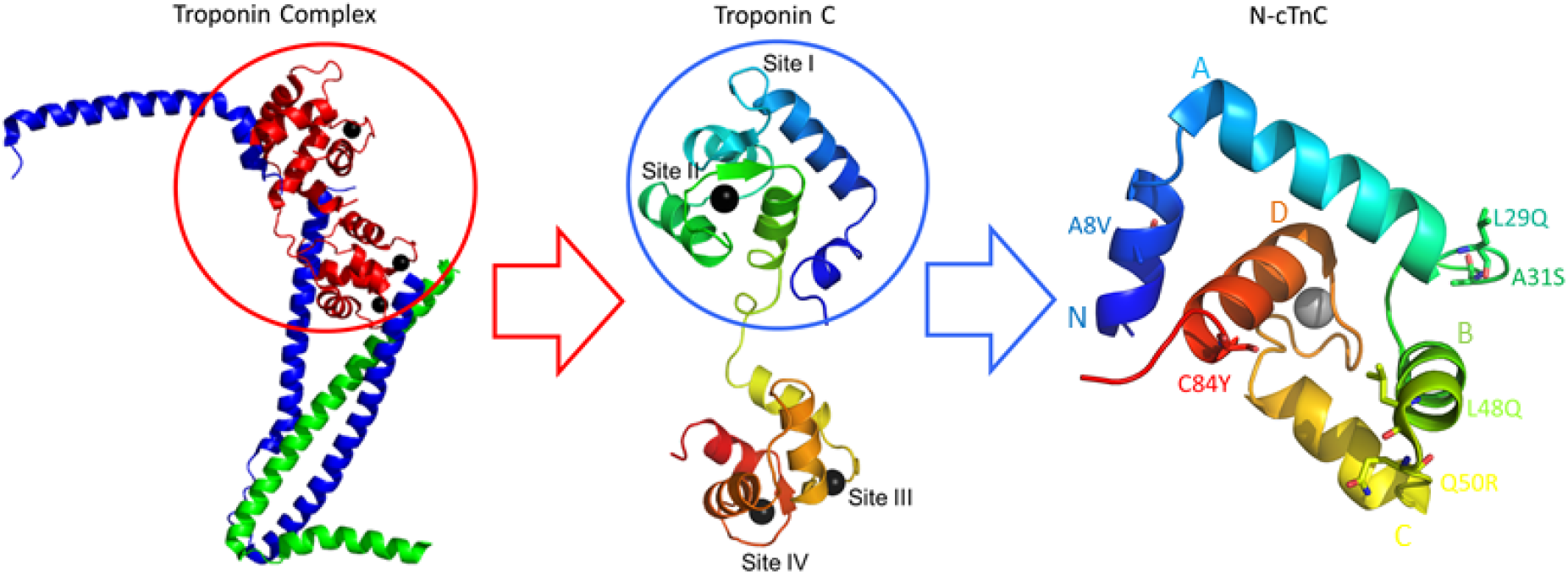
N-cTnC within cTnC and the Troponin complex, the cTn complex is shown on the left panel. This complex includes the Ca^2+^ binding cTnC in red, the inhibitory cTnI in blue, and the tropomyosin binding cTnT in green. Black spheres depict the bound cations that interact with sites III/IV in the C-terminal domain and site II in the N-terminal domain. cTnC, shown in the middle panel in rainbow colors with the N-terminal domain in blue and the C-terminal domain in red. The black spheres again depict the bound cations. The focus of this work is on the N-domain of cTnC which is depicted in the right-most panel with each of the 6 studied mutations highlighted and labelled. In the right most panel, each of the helices within N-cTnC is labelled and color coded to correspond to depicted structure. From the N terminus, these are helices N through D.

Hypertrophic Cardiomyopathy (HCM) afflicts ~1 in 200 in the general population (7,8). Over 1,000 HCM-associated mutations have been found in a variety of sarcomeric proteins, of which over 100 are located in the cardiac troponin complex (9–11). Despite a wide range of molecular precursors, the disease phenotype consistently entails some degree of hypertrophy, myocyte disarray, and fibrosis resulting from a wide variety of underlying causes (12,13). This devastating disease often manifests as sudden cardiac arrest secondary to ventricular tachycadia/fibrillation and is the most common cause of sudden cardiac death in young athletes (14);(15).

Increased Ca^2+^ sensitivity causes a prolongation of the contractile phase and a reduction in relaxation, which is thought to elicit hypertrophy (4). The role of cTnC as the Ca^2+^-sensing component has made it a target of study with multiple HCM-associated mutations identified in the regulatory N-terminal domain (**Figure 1**)(16,17). Here, we focus on A8V (18), L29Q (19), A31S (20), and C84Y (21), the engineered mutation L48Q (3) and the Dilated Cardiomyopathy (DCM) associated mutation Q50R (22) (**Figure 1**). We have previously studied this series of mutations through Isothermal Titration Calorimetry (ITC) and Molecular Dynamics (MD) Simulations (23). Our findings support the notion that mutations which destabilize the closed-state of cTnC, or those that stabilize the interaction with the TnI_SW_ in the open-state increase the Ca^2+^-binding affinity, as was the case with L48Q, Q50R, and C84Y but not A8V, L29Q, A31S.

In this study, we have chosen to utilize ITC, avoiding any interference from fluorescent reporters (24–34).

After potassium, Mg^2+^ is the second most abundant cellular cation with a total concentration of ~15 - 20 mM. Mg^2+^ is also tightly regulated through extensive buffering by cytosolic components such as ATP. Even still, free Mg^2+^ measures approximately 0.5 - 1 mM (though a wide range of numbers between 0.2 – 3.5 mM have been reported in different systems) and is about 1,000-fold more abundant than systolic free Ca^2+^ (35–37). As the majority of cellular Mg^2+^ exists in complex with ATP, limited ischemia (15 mins) elevates free Mg^2+^ up to 10-fold (to 6.5 mM) and reperfusion (30 mins) restores Mg^2+^ back to near baseline levels (38,39) because of ATP depletion and the replenishment. Studies on isolated TnC (40–45), the cTn complex (40,42), and reconstituted fibers (42,46) have demonstrated that Mg^2+^ decreased the ability of Ca^2+^ to induce structural change. Previous studies utilizing physiological systems similarly demonstrated an inverse correlation between Ca^2+^ sensitivity of force production and availability of Mg^2+^ in skinned skeletal and cardiac fibers (47–49). 3 mM Mg^2+^ decreases the Ca^2+^ affinity in isolated cTnC (3) and skinned psoas muscles (27), but does not seem to cause confomational changes in cTnC (3).

A prevalent hypothesis posits that mutations which destabilized the closed conformation of the protein prior to Ca^2+^ binding and/or those that favor the open, Ca^2+^-bound state confer an increase in affinity (50). Sequence variations outside the coordinating residues of the EF hands of cTnC may induce alterations in Ca^2+^ affinity allosterically (51,52). These changes in Ca^2+^ binding have previously been linked to HCM and DCM-associated mutations, thus insight regarding the underlying thermodynamics of this foundational interaction may have far reaching implications (23,28,53). In contrast, mutations outside the binding residues of each EF hand are not thought to allosterically modify Mg^2+^ binding (54–56), therefore the role of this cation in HCM is currently unclear. Here, we further explore the effects of Mg^2+^ on Ca^2+^ binding to the regulatory domain of cTnC and possible modifying affects on the previously listed series of mutations.

Previous ITC and thermodynamic integration (TI) simulations corroborate these findings; Mg^2+^ competes with Ca^2+^ for binding to site II and at physiological concentrations (1 mM) significantly reduces Ca^2+^-binding in full-length and N-terminal cTnC (57). In order to obtain a more complete picture of the cardiomyopathy mutations, it is important to establish whether they affect binding of Ca^2+^, Mg^2+^, or both. If the binding of these ions is affected differentially by a specific mutation, then this could exacerbate or attenuate the effect when considering Ca^2+^ alone. We observed that the change in Ca^2+^ and Mg^2+^ binding affinity was not the same for the each construct. In particular, the affect of L48Q, Q50R, and C84Y on the binding of these cations was significantly different from the WT and underlines the importance of considering background Mg^2+^ levels which were studied through competition experiments. Here, we show that the affinity of five HCM associated variants and one DCM associated variant for Mg^2+^ was significantly higher than WT cTnC, each was also found to have a physiological significant affinity for Mg^2+^. Therefore, the presence of baseline Mg^2+^ may contribute to the dynamics which govern cardiac excitation-contraction coupling. Further, higher than baseline concentrations of Mg^2+^ that accompany energy-depleted states causing ischemia may accentuate the effects of inherited mutations and further exacerbate dysfunction of the diseased heart.

## Results

For each construct: WT, A8V, L29Q, A31S, L48Q, Q50R, and C84Y, Ca^2+^ and Mg^2+^ were independently titrated into apo-state N-cTnC out to measure the affinity for binding of each cation as discussed in the Methods section (**Figure 2**). Subsequently, each construct was pre-incubated with 1 mM Mg^2+^ and 3 mM Mg^2+^ independently and titrated by Ca^2+^ to measure the relative binding affinity of Ca^2+^ in these conditions (**Figure 3**).

**Figure 2:**
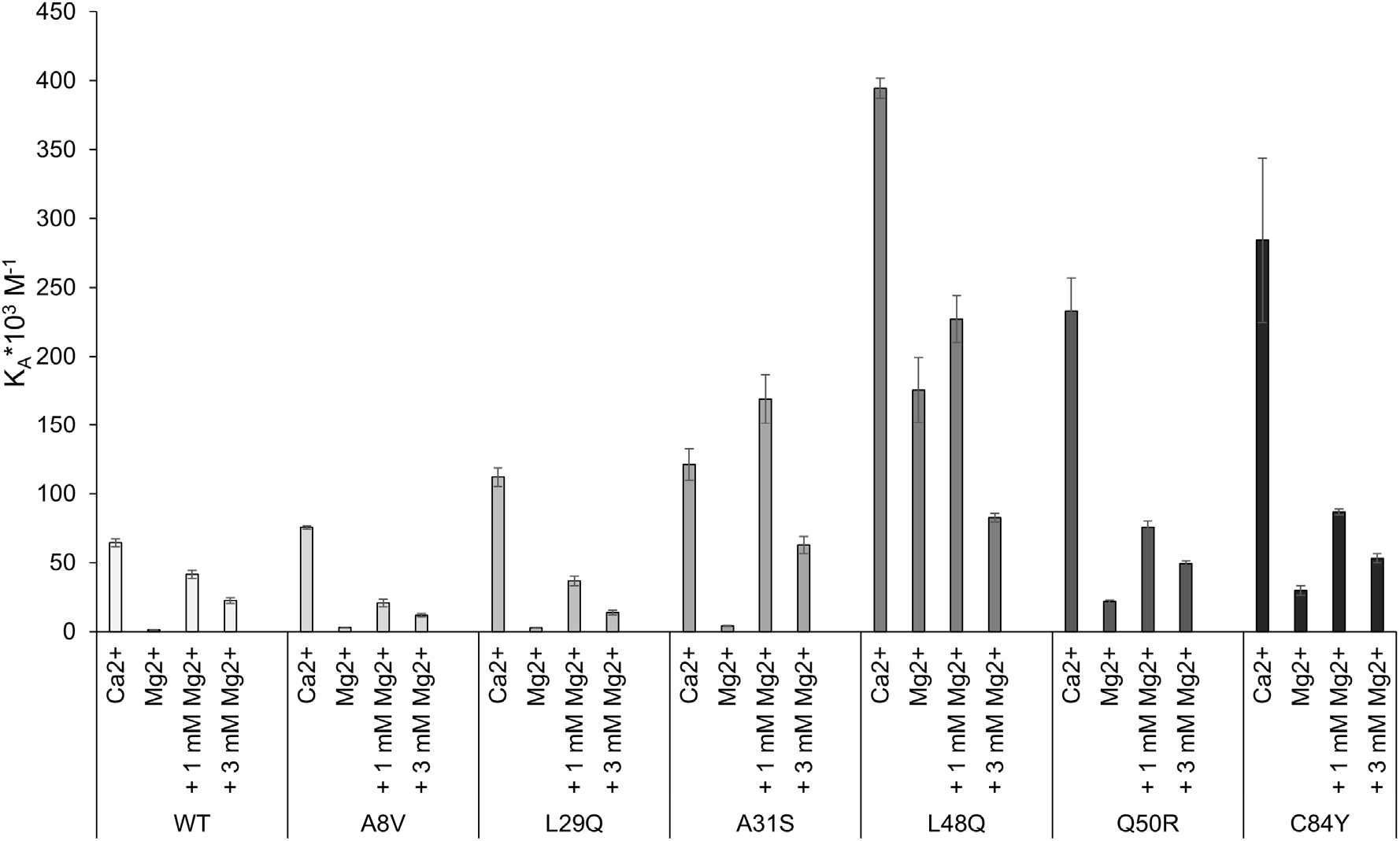
Comparing the affinity for Ca^2+^/Mg^2+^ in each titration condition between all N-cTnC constructs. SEM error bars are used to depict where significant differences exist in the mean values. With the exception of L48Q, the highest affinity is seen in the Ca^2+^ titration and the lowest in the Mg^2+^ titrations. L48Q has the highest Mg^2+^ binding affinity by over an order of magnitude. Increasing Mg^2+^ from 0 to 1 to 3 mM lowered Ca^2+^ binding affinity in a graded manner. The more C-terminal mutations cause a greater increase in Ka relative to the WT.

**Figure 3.**
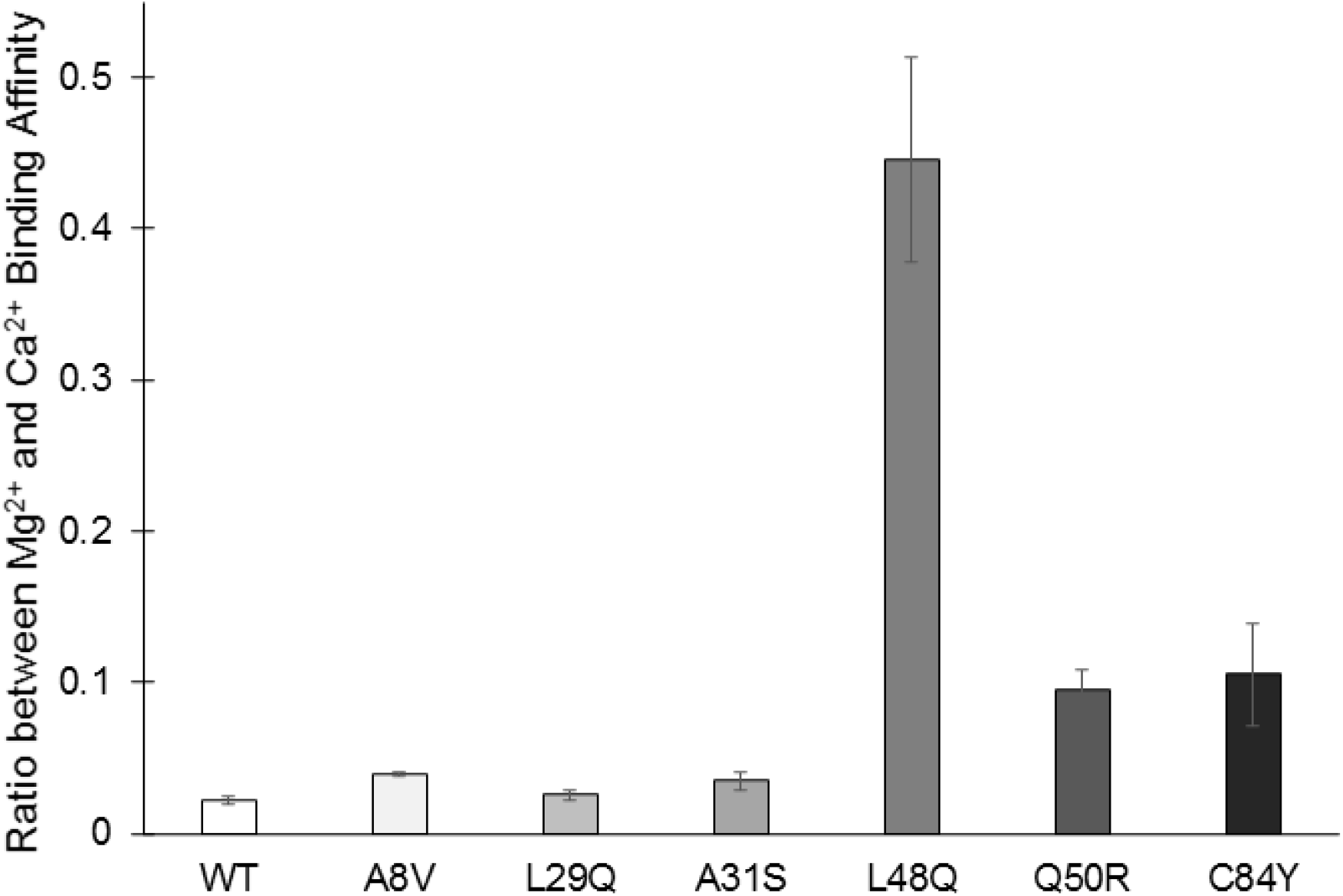
Ratio between Mg^2+^ and Ca^2+^ binding affinity for each N-cTnC construct indicating the realtive change in affinity. Error bars are also depicted to provide a measure of the confidence in each ratio. A greater ratio indicates that the Mg^2+^-binding affinity was less reduced in the WT/mutant construct relative to the baseline Ca^2+^-binding affinity. The ratio is not equal to one in any of the constructs highlighting the importance of considering background cellular Mg^2+^ in these studies and stressing the importance of competition experiments carried out here.

### Construct Based Comparison

In the WT N-cTnC and all of the mutants, the presence of Mg^2+^ significantly altered the Ca^2+^ binding affinity. The mutants all had a significantly lower K_d_ in the presence of 1 mM Mg^2+^ in comparison to WT N-cTnC. In each of these mutants, Ca^2+^ binding to site II occurred with significantly lower affinity in the presence of Mg^2+^.

#### WT N-cTnC

The interaction of Ca^2+^ with WT N-cTnC was endothermic and entropically driven. The K_d_ value associated with the interaction was 15.79 ± 0.74 μM. Mg^2+^ affinity was more than an order of magnitude lower but similarly endothermic and entropically driven. 1 mM Mg^2+^ lowered binding affinity 1.6-fold and reduced the amount of binding (as indicated by lower ΔH) and the entropic favourability of the interaction. 3 mM Mg^2+^ further accentuated these changes. These results are in agreement with those obtained previously (23,57).

#### A8V N-cTnC

A8V exhibited isotherms that were similar to WT with the binding affinity being only moderately higher for both Ca^2+^ and Mg^2+^. Both interactions were endothermic and entropically driven. The addition of 1 mM Mg^2+^ reduced the Ca^2+^ affinity approximately 3-fold and significantly reduced the amount of binding to site II (ΔH was reduced to less than half). The effects of 3 mM Mg^2+^ pre-incubation was similar in that the KA, ΔH, and ΔS were reduced such that the reactions were less spontaneous, and binding occurred with a K_d_ that was nearly 7-fold higher than the apo-state condition. Therefore, the effect of Mg^2+^ pre-incubation on Ca^2+^ binding was more significant in the A8V mutant compared to the WT protein.

#### L29Q N-cTnC

The changes observed in L29Q relative to the WT protein was in line with the trends seen in the previous mutant. The affinity was higher than seen in the WT protein and the interaction was again, endothermic and entropically driven. The affinity of the mutant was nearly 2-fold higher than seen in the WT. The addition of 1 and 3 mM Mg^2+^ decreased the affinity such that K_d_ increased nearly 3- and 9-fold, respectively. The ΔH, indicative of the amount of binding also decreased in a graded manner with increasing amounts of Mg^2+^ such that a favorable ΔS was nearly the sole factor driving this interaction.

#### A31S N-cTnC

Ca^2+^ binding to A31S was endothermic and entropically driven and occurred with higher affinity than seen in the WT protein. The affinity of the mutant for Mg^2+^ was 3-fold higher than the WT. Given this, it was not surprising that the addition of 3 mM Mg^2+^ decreased the affinity for Ca^2+^ to about half of that seen in the apo-state binding condition. However, two surprising findings were seen. Firstly, the addition of Mg^2+^ altered the reaction kinetics such that the interaction became exothermic. Secondly, the 1 mM preincubation condition was characterized by higher than apo-state affinity for Ca^2+^. This may be due to the more than 30-fold higher affinity for Ca^2+^ such that 1 mM Mg^2+^ enhanced binding, despite assumed occupation of some percentage of the binding sites. Even still, the affinity for Ca^2+^ was further reduced when Mg^2+^ was elevated to 3 mM.

#### L48Q N-cTnC

Among the constructs studied, L48Q had the highest affinity for both Ca^2+^ and Mg^2+^ by a significant and substantial margin. It was characterized by an exothermic interaction when binding Ca^2+^ and an endothermic interaction with Mg^2+^. The addition of 1 mM Mg^2+^ decreased the affinity by less than 2-fold and 3 mM Mg^2+^ decreased the affinity by nearly 5-fold. In both pre-incubation conditions, the interaction with Ca^2+^ was still exothermic, but the change in entropy was negligibly small such that the reaction was driven almost entirely by the dissipated heat.

#### Q50R N-cTnC

The affinity of Q50R for both Ca^2+^ and Mg^2+^ was higher than the WT. Binding was similarly endothermic and characterized by a positive ΔS. However, the addition of 1 mM Mg^2+^ had significant effects on the reaction such that binding of Ca^2+^ occurred with one third of the affinity and the interaction became exothermically driven as the change in entropy was much less favorable (about one quarter the T*ΔS). The addition of 3 mM Mg^2+^ further reduced the affinity such that it was about 5-fold lower but still exothermic. It should be noted that with a greater concentration of pre-incubated Mg^2+^, the amount of binding was reduced as seen through a lower ΔH.

#### C84Y N-cTnC

After the L48Q mutation, C84Y caused the largest change in affinity for both Ca^2+^ and Mg^2+^ relative to the WT protein. This was in keeping with the trend of the C-terminal most mutations having a larger effect and enacting more significant changes on the energetics of these titrations. In the apo-state, the interaction with both cations was endothermic (although the absolute value indicated that the amount of binding was much less than the WT) and entropically driven. The relative contribution of these parameters indicates that entropic favorability played a relatively large role in driving these reactions. Preincubation with 1 mM and 3 mM Mg^2+^ reduced affinity by approximately 3 and 5-fold respectively, with the reaction becoming exothermic.

### Disease based comparison

Only one DCM-associated mutation (Q50R) was studied in this series of experiments. Details regarding this construct are discussed above. In brief this construct had a greater than WT binding affinity for both Ca^2+^ and Mg^2+^, thus it was sensitizing to both in our experiments. The effect on Mg^2+^ binding, relative to the baseline Ca^2+^ binding was less than seen in the WT (**Figure 3**). In the competition experiments, a significant change in thermodynamics was seen, whereby the interactions become enthalpy driven and exothermic in nature. Consistent with higher overall affinity, in the preincubation experiments the reduction in Ca^2+^ binding affinity was less than seen in WT N-cTnC. All other mutants were HCM-associated, though L48Q is an engineered mutation. Each construct had a higher than WT Ca^2+^-binding affinity and with the exception of A8V/L29Q, each had a higher than WT Mg^2+^-binding affinity (**Figure S1**). There does not seem to be a clear distinction in pattern between the DCM/HCM constructs in this system.

### Titration based comparison

#### Calcium binding

L48Q, Q50R, and C84Y had significantly higher affinities than WT (**Figure S1**). L29Q and A31S also changed the K_d_ significantly compared to the WT protein with the A8V mutation being the only construct that did not cause a significant elevation in affinity for Ca^2+^. L48Q, Q50R, and C84Y had lower than WT changes in entropy. The Ca^2+^-L48Q interaction was the only exothermic titration.

The ΔG reflects both the ΔH and the ΔS and as such demonstrates that the most favorable interaction occurs in L48Q, then Q50R/C84Y, followed by the other mutants.

#### Magnesium Binding

It is important to note that the interactions with Mg^2+^ were endothermic and entropically driven for all 6 mutants and the WT protein (**Table 2**). The highest affinities were seen in L48Q, Q50R, and C84Y. While the Mg^2+^ K_d_ was significantly lower for all mutants compared to WT, with L48Q being the lowest by two orders of magnitude. The relative difference in Mg^2+^ and Ca^2+^ binding affinity shows patterns with the L48Q>>C84Y~Q50R compared to the WT (**Figure 3**). These mutants affected Mg^2+^ and Ca^2+^-binding to a different degree, increasing Mg^2+^-binding more than they increased Ca^2+^-binding. In the presence of competing Mg^2+^ in the cellular milieu, a greater effect on Ca^2+^-binding would be expected and they may be termed: “more severe”. With the exception of L48Q, the ΔH values for each mutant and WT construct were very similar and the elevated Mg^2+^-affinity was due to higher ΔS values. The exception is L48Q in which a significantly greater amount of binding occurs as indicated by the absolute value of ΔH. Every mutant was associated with a higher ΔS than the WT, with the highest values being associated with L48Q, followed by Q50R, and subsequently C84Y.

#### Competition

To gain a measure of the affinity of Ca^2+^ for each construct in the presence of competing/background Mg^2+^, it is possible to look at the ratio between titrations (**Figure 3**). A number of the constructs responded similarly to the WT to each of Ca^2+^ and Mg^2+^ with the exception of: L48Q, Q50R, and C84Y. The ratio between the Mg^2+^ and Ca^2+^ binding affinity of these mutants was significantly greater than the WT construct. These findings stress the need for competition experiments as the Ca^2+^-affinities should be studied in the presence of physiologically relevant Mg^2+^ concentrations.

### 1 mM Magnesium Competition

As expected from the affinity for each individual cation, the L48Q construct had the highest affinity for Ca^2+^ in the 1 mM Mg^2+^ pre-incubation condition. Significant increases in Ca^2+^ affinity were also observed for A31S, Q50R, and C84Y, with A8V and L29Q not distinguishable from the WT (**Table S5**).

The amount of binding can be tracked through the value of ΔH. The greatest change was seen in A8V, L29Q, A31S, and L48Q that are also driven by exothermic interactions. The ΔH values for A8V and L29Q, while different from WT, were nonetheless similarly endothermic.

The same four mutations: A31S, L48Q, Q50R, and C84Y showed a less favorable entropy change compared to the WT. Despite this, the binding was more spontaneous and was driven by the change in ΔH associated with each interaction. A8V and L29Q also interact spontaneously due to a favorable change in ΔS.

### 3 mM Magnesium Competition

The patterns observed in the 1 mM pre-incubation condition were largely maintained and accentuated further in the 3 mM pre-incubation condition. In general, the highest KA was seen in the mutants adjacent to site II, with A8V and L29Q statistically indistinguishable from the WT construct (**Table S6**).

L48Q had the highest affinity for Ca^2+^ in the presence of 3 mM Mg^2+^ as expected from its affinity for each individual cation. The ΔH was characteristic of an exothermic interaction in the A31S, L48Q, Q50R, and C84Y constructs and drove these more spontaneous reactions. All conditions had a positive ΔS, but the contribution was most favorable for WT, followed by L29Q, A8V, A31S, Q50R, C84Y, and L48Q which had the smallest ΔS contribution. A8V and L29Q are most WT like; they have the most favourable changes in entropy that resulted in spontaneous interactions despite endothermic binding.

### Mg^2+^ binding affinities from Thermodynamic Integration

Thermodynamic Integration (TI) was conducted to calculate the binding affinity (Δ*G_TI_*) for Mg^2+^ to WT N-cTnC, L48Q N-cTnC, Q50R N-cTnC, and C84Y N-cTnC. The binding affinity for each protein structure was averaged over 5 independent runs and was −5.139 ± 2.308 *kcal/mol*, −5.481 ± 0.719 *kcal/mol*, −6.205 ± 2.112 *kcal/mol*, −6.364 ± 1.372 *kcal/mol*, respectively. The TI calculated Mg^2+^ binding affinities were in good agreement with the ITC data, except for L48Q N-cTnC. The ITC data showed a much stronger increase in Mg^2+^ sensitivity for the L48Q N-cTnC. However, all mutated structures were shown to have increased Mg^2+^ sensitivity compared to the wild type structure. The ΔΔ*G*_*Q*50*R*_ values were similar for TI and ITC (1.066 *kcal/mol* and 1.630 *kcal/mol*, respectively). The ΔΔ*G*_*C*84*Y*_ values also showed good agreement for TI and ITC (1.225 *kcal/mol* and 1.790 *kcal/mol*, respectively) (**Table 1**).

**Table 1.** Mg^2+^ thermodynamic integration free energy of WT and mutant N-cTnC structures

| N-cTnC<br>Structure | $\Delta G_{TI}$ (kcal/mol)<br>± std. dev. | $\Delta\Delta G_{TI}$ |
| --- | --- | --- |
| WT | $-5.139 \pm 2.308$ | |
| L48Q | $-5.481 \pm 0.719$ | 0.342 |
| Q50R | $-6.205 \pm 2.112$ | 1.066 |
| C84Y | $-6.364 \pm 1.372$ | 1.225 |

### Summary of results

All of the studied mutations increased Ca^2+^ and Mg^2+^ binding affinity relative to the WT construct (**Tables 1 and 2**). The smallest increase was associated with A8V and the greatest with L48Q. In general, the mutations adjacent to site II (L48Q, Q50R, and C84Y) increased the affinity for each cation to a greater extent and altered the thermodynamic profile of the binding interaction more dramatically; the changes associated with the other mutants were more subtle.

The K_A_ associated with Mg^2+^ binding was orders of magnitude lower than that with Ca^2+^ binding for all constructs (**Figure 2 and S5**). The addition of 1 mM Mg^2+^ decreased the affinity for Ca^2+^ but also decreased the extent to which Ca^2+^ was able to bind site II (as visualized through a reduction in ΔH) (**Table S5**). Addition of 3 mM Mg^2+^ further decreased both the K_A_ and ΔH (**Table S6**).

Mutations with the closest proximity to site II, namely Q50R and C84Y responded most dramatically to the presence of Mg^2+^. Pre-incubation significantly altered the reaction landscape such that endothermic apo-state interactions became exothermic.

## Discussion

The binding of Ca^2+^ to site II within the N-terminal domain of cTnC is the fundamental molecular precursor to a series of conformational changes that culminate in force production. As such, changes in the sequence of this highly conserved protein often have grave consequences for the force production capabilities of the heart (58) which likely lead to cardiac remodeling. The 6 mutations that we have studied are all situated outside the EF hand binding regions and allosterically alter Ca^2+^ affinity. Given the location of each mutation of interest and the desire to focus on changes in the binding interaction, we exclusively studied the N-terminal domain of cTnC. The mutations in question have also been studied at various levels of complexity by numerous groups, whose findings are in general agreement with work done by our group and others (3, 18–23).

The recently published Cryo-EM structure of the cardiac thin filament has shown that cardiomyopathy associated mutations in troponin overwhelmingly occur in regions that interface with the actin-tropomyosin complex (59). Mutations which occur at a distance from these interfaces, are still most likely to affect changes through altered interactions with other proteins of the contractile complex (60). Loss-of-function mutations in troponin increase the Ca^2+^ sensitity of the contractile element as they decrease steric hinderance of myosin binding sites. In contrast, gain-of-function mutations in N-cTnC have a sensitizing effect given that in the holo-state, this domain displaces the TnI_SW_ from actin, moves tropomyosin, and unhinders myosin binding sites (61). However, very few pathogenic mutations have been identified in cTnC, indicating lethality consistent with the critical role of this molecule.

ITC directly measures the binding interaction and subsequent conformational change as an alternative to the introduction of naturally occurring fluorophores such as F27W (24) or synthetic fluorophores such as IAANS (28,62). These reporters can be used to quantify the structural changes that proceed the binding interaction and thus, indirectly measure affinity. Fluorescence can also be used to report on the dissociation of Ca^2+^ from cTnC, cTn, or the TF by utilizing chelators such as EDTA and rapid fluid changes through stopped flow experiments (52).

Equilibrium dialysis studies suggested that Mg^2+^ does not compete with Ca^2+^ for binding to the N-terminal cTnC, and exclusively binds the C-terminal sites (63). This notion has endured for decades. Since then, several studies have shown through a number of different experimental techniques that the Mg^2+^ binding affinity of site II of skeletal and cardiac TnC is physiologically relevant (41,43,49,64–70). The Mg^2+^ affinity of site II has was estimated through fluorescence to have a K_d_ of ~1.5 mM and through competition experiments, site II was posited to be 33 – 44% saturated with this cation at diastolic concentrations of Ca^2+^ (3). More recent studies show that these results also hold in TnC variants of similar sequence from other species where the binding affinity of Mg^2+^ is an order of magnitude lower than Ca^2+^ (71). Our recently published ITC and thermodynamic integration simulations corroborated these findings (57).

Our studies utilize ITC, that has the sensitivity to detect minute enthalpic fluctuations which accompany these interactions, accurately detecting changes to within 0.1 μcal (34). That is, not withstanding the inherent limitations which are part and parcel of each experimental technique. For this assay a limitation results from the absence of other components of the cTn complex, particularly cTnI which plays a central role to the regulation of TF Ca^2+^ handling (28,72,73); thus care must be taken when translating these findings to more complex systems.

The binding of Ca^2+^ to N-terminal cTnC is driven through a balance between the conformational strain resulting from the interaction and the energetics of exposing a hydrophobic cleft to the aqueous environment (74). We posit that the changes in affinity seen in each mutant result from either the destabilization of the apo-state protein or the stabilization of the solvent exposed state (23).

In general, we found that the mutants studied all had a more negative ΔG compared to the WT, consistent with our previously published MD Simulations (23). Work by Bowman and Lindert corroborates these findings and suggests a unifying theory that increased frequency of opening may result from the lowered energetic cost of exposing the N-terminal domain of cTnC. The placement of more hydrophilic amino acids that destabilize hydrophobic packing in the closed state and stabilize the open, solvent-exposed state that follows, may allow for this mechanism of action (75).

A8V was only moderately different from the WT, consistent with previous findings that suggest this mutation alters the interaction of cTnC with other TF proteins rather than altering the Ca^2+^ binding affinity directly (21,72). The locus of this valine near the interface with N-cTnI and strengthened interaction with the switch peptide makes this a distinct possibility (23,76). Nuclear Magnetic Resonance (NMR) data suggest a slight increase in the opening frequency in the apo-state relative to the WT (18). In our study, the binding affinity of this mutant for Mg^2+^ was higher than the WT and as a result, in the pre-incubation conditions, the A8V construct had lower than WT affinity for Ca^2+^ (**Figure 6 and Table 6**).

L29Q is the next most proximal mutation to the N-helix and caused a statistically significant reduction in the K_d_ for Ca^2+^ binding compared to the WT. It also had greater than WT Mg^2+^ binding affinity but similar thermodynamic parameters and isotherm characteristics. Fluorescence studies on isolated cTnC containing the L29Q mutations showed increased Ca^2+^ affinity (28). A complex system containing the entire contractile apparatus with L29Q cTnC had similar to WT Ca^2+^ sensitivity (77). This mutation also changes sensitivity of force generation in a length and phosphorylation-dependent manner (51).

Alteration of a hydrophobic residue to one that is polar underlies sensitization of site II (78,79). Mechanistically, solvent exposure of an uncharged glutamine may facilitate a greater extent of opening than a hydrophobic leucine (23). However, our previous work suggests that this mutant has the highest closed probability amongst the 7 constructs and lowers opening frequency (23). In contrast, it has been shown through NMR that this mutation may cause a more open N-domain in the cTnC in both the apo- and holo-states (80). L29Q may open in the holo-state with similar frequency as the WT and have similar energetic requirements as the WT for opening in both the apo- and holo-states (75). Therefore, it is likely that the effects of this mutation in isolated N-cTnC are minimal and that changes are enacted through modification of the interaction with other cTn complex proteins (28).

A31S is in the EF hand of the defunct site I and changes a hydrophobic amino acid for an uncharged one. In skeletal tissue, there is a great deal of cooperativity between binding to the two N-terminal sites of TnC (81). This mutant had a significantly lower K_d_ for both Ca^2+^ and Mg^2+^ compared to the WT and significantly higher affinity for Ca^2+^ in both pre-incubation conditions. A31S was found through MD simulations to sample a greater number of interhelical angles and to have a lower average angle between helices A and B (23). The most interesting finding was that pre-incubation with 1 or 3 mM Mg^2+^ completely changed the reaction isotherm (**Figure S3**).

ΔH reflects the strength of hydrogen bonds, van der Waals interactions, and electrostatic forces between the titrant and the target ligand. Optimal placement of hydrogen bond donors and acceptors balances de-solvation of polar groups to contribute to the enthalpy change (82). The significant change in enthalpy suggests alteration of the number of bonds formed by the side chains of binding site residues or those exposed to the environment following the conformational change. This mutation may stabilize the binding site I between helices A and B through formation of an additional hydrogen bond causing local changes that minimally alter global structure (20).

L48Q caused by far the largest change in affinity and altered the thermodynamics of each isotherm to the greatest degree. This mutation is located within the BC helical bundle and was strategically engineered to increase the Ca^2+^ sensitivity of force production (3). The combination of high Ca^2+^ and Mg^2+^ affinity resulted in the lowest observed K_d_ in both competition conditions.

Our previously published MD simulations suggest that the absence of a hydrophobic residue disrupts hydrophobic packing in the AB domain. We also found that the L48Q mutant opens more frequently than the other constructs (23). Bowman and Lindert’s work also suggests that L48Q samples the open state much more frequently than the WT protein (75). The changes in ΔH are likely, at least in part, due to the presence of an additional hydrogen bond resulting from the introduction of a polar amino acid in a key domain of cTnC. *In vivo*, this would increase the opportunities for interaction with the TnI_sw_ (15,83,84).

Tikunova et al. originally suggested that despite a shift towards the Ca^2+^-bound state, resulting from a reduction in hydrophobic contact between helices NAD and BC, the solvent exposure of the N-domain is minimized by numerous side chain contacts. Their hypothesis regarding the disruption of hydrophobic interactions and minimization of exposure to the surrounding solvent is reconcilable with our findings and explains the much lower ΔS associated with this set of titrations (3).

Q50R is a relatively recently identified mutation that has yet to be fully explored. This change replaces a polar side chain with one that is bulkier and charged. Given the hypothesized form of action seen in L48Q and the vicinity of these residues, it is conceivable that the packing of helices NAD and BC is also disrupted by this mutant. This mutant had a much higher affinity than WT for both Ca^2+^ and Mg^2+^ which bind endothermically. Similar to L48Q, the interaction with Ca^2+^ was exothermic in the pre-incubation condition. Our previous work suggests that Q50R is more frequently open than the WT cTnC (23). The reduced entropic cost of exposing a charged residue to the aqueous environment may explain the decreased ΔS of the system. Further, the energetic cost of opening the hydrophobic patch is increased in this mutant in comparison to the WT protein that has a less stable closed conformation (75).

C84Y places a bulky hydrophobic side chain in the region immediately preceding the flexible DE linker that is bound and stabilized in the open state by the TnI_SW_. This bulky tyrosine may act as a wedge to reduce interaction with the TnI_SW_ and thus increase the Ca^2+^ sensitivity of force development in skinned fibers (21,72). This mutation was thermodynamically similar across all titrations with Q50R, which given the location of each does not necessarily suggest a similar mode of action. Interestingly however, our MD Simulations previously showed that a hydrophobic interaction between C84 and Q50 may be disrupted by this mutation. The bulky tyrosine in helix D may reduce the entropic cost of opening associated with the binding interaction, this is consistent with the observed, lower than WT ΔS values in C84Y N-cTnC (23).

The calculated TI Mg^2+^ binding affinity results were in good agreement with the ITC values. We observed increased ion sensitization for all mutant N-cTnC structures. In particular the ΔΔ*G_TI_* values for Q50R and C84Y mutations aligned very well with the experimental data. The absolute binding affinities calculated for WT, Q50R and C84Y were overestimated compared to the ITC values by less than 1 kcal/mol. Parameterization of cations, especially Mg^2+^, in simulating biological systems has proven to be difficult (85,86). This could offer a potential explanation for the overestimation and relatively high standard deviations observed in the averaged absolute binding affinities. Another source of discrepancy between the *in silico* and ITC results, arises from the fact that there is no PDB structure of Mg^2+^ bound to site II of N-cTnC. In order to create the starting structure for TI simulations the PDB 1AP4 served as the base model, and Mg^2+^ was substituted in place of the Ca^2+^ ion. If crystal structures of Mg^2+^-bound WT N-cTnC and mutants existed, the use of these structures could potentially improve the IT results. TI of the L48Q mutant did not produce nearly as strong an increased Mg^2+^ sensitization as observed in the ITC data. We speculate that this could possibly be attributed to large conformational changes that were unable to be captured using TI. The timescale of the TI simulations was only 5 ns, which was insufficient to properly sample any large protein conformational changes.

Our results here show that WT N-cTnC and each of the mutants responded significantly and variably to the presence of Mg^2+^. Except for A31S, each mutant site II has a significantly lower Ca^2+^ binding affinity in the presence of 1 mM Mg^2+^. Desensitization was most pronounced in A8V>L29Q>Q50R>C84Y>L48Q. Given these observations, it is possible that Mg^2+^ binding dampens the presupposed sensitizing effect of HCM-associated mutations (Chang and Potter 2005); at the very least, the effect of background Mg^2+^ in modifying Ca^2+^ sensitivity of force production cannot be ignored (**Figure 2**).

Given the high concentration of free Mg^2+^ in the cytosol, its similarities as a divalent cation and the small difference in atomic radius with Ca^2+^, suggests that this ion is a candidate for binding to site II of cTnC. Despite previous work in this field, the central dogma in the literature is largely dismissive of the possibility that physiologically relevant concentrations of Mg^2+^ bind to site II. A polar serine at residue 69 and a negatively charged glutamic acid at residue 76 in the EF hand binding site II of N-cTnC, create a domain that is conducive to Mg^2+^ binding (87,88). We previously established the binding of Mg^2+^ to site II in full length and N-terminal cTnC and would compete with Ca^2+^ for binding to this site at physiological concentrations of each cation in the cell (57). In this work, we explore the hypothesis that Ca^2+^ and Mg^2+^ compete for binding, where affinity for each cation is allosterically modified by single amino acid changes outside the binding domain.

Tikunova and Davis have shown that Mg^2+^, unlike Ca^2+^, does not cause a structural change upon binding but does significantly alter the affinity of cTnC for Ca^2+^ with 3 mM Mg^2+^ causing a more than 3-fold reduction in Ca^2+^ binding affinity. Moreover, Mg^2+^ reversed the fluorescence change of Ca^2+^ saturated cTnC; that is to say, 3 mM Mg^2+^ competes for binding to site II (3).

The molecular mechanisms which underpin the role of cellular Mg^2+^ in cardiac contractility have yet to be fully understood and require further exploration. We measured a 47-fold difference in the affinity of WT N-cTnC for Ca^2+^ (15.79 ± 0.74 μM) in comparison to Mg^2+^ (711.08 ± 32.67 μM). However, Mg^2+^ is at least 1000 times more abundant in the cytosol at systole than Ca^2+^ 1 μM vs. 1 mM (89,90) and may compete for binding to site II in addition to the structural sites III and IV (57). Mg^2+^ deficiency has been linked to cardiac disease including arrhythmias, hypertension, and congestive heart failure (91–94). It is possible that Mg^2+^ modulates the role of Ca^2+^ and alters activation of contractile pathways that are governed by this messenger.

Less than 15 mins of ischemia can decrease [ATP]i resulting in an increase in free [Mg^2+^] by three-fold (95). This elevated Mg^2+^ may compete with Ca^2+^ for binding to cytosolic buffers. Overall, our study suggests that the effect of cellular Mg^2+^ on the Ca^2+^ binding properties of site II within N-cTnC is not negligible. This effect may be even more pronounced in HCM- and DCM-mutant N-cTnC, where both cytosolic concentrations of free Mg^2+^ (1 mM) and elevated Mg^2+^ that may accompany energy depleted states (3 mM) caused a more significant reduction in affinity compared to the WT through alterations in structural dynamics and the energetic landscape of each binding interaction.

The limitations of these studies at the levels of molecular complexity undertaken here are clear and have been outlined above. Despite these shortcomings, the utility of a detailed understanding of the molecular underpinings of disease entities such as HCM is essential for progress towards the age of personalized medicine. This understanding is can be achieved through myriad techniques that are increasingly utilized (13). To date, treatment of HCM has largely focused on symptom management which in some patients has limited efficacy. A precision medicine based approach has been proposed instead, by which patients are subdivided by underlying mechanism and treated accordingly (60).

## Conclusions

The interaction of Ca^2+^ with mutant N-cTnC occurred with higher than WT affinities, with the highest affinity seen in the L48Q mutant. In general, A31S, L48Q, Q50R, and C84Y had the highest affinities for both Ca^2+^ and Mg^2+^. Thermodynamic, structural, and simulation work by our group and others suggests a common mechanism whereby mutants destabilize hydrophobic interactions between helices NAD and BC cause an elevation in binding affinity.

We found that the affinity for Mg^2+^ was at least an order of magnitude lower than seen for Ca^2+^. The change in affinity observed when comparing the Mg^2+^ pre-incubated N-cTnC and apo-state protein was variable in each mutant and significantly different from the WT. Moreover, 1 mM and 3 mM Mg^2+^ caused a graded decrease in the amount of binding and affinity for Ca^2+^. In contrast to Ca^2+^, cellular Mg^2+^ does not cause a conformational change upon binding to site II of cTnC and thus cannot initiate contraction. However, Mg^2+^ has been shown, both here and in numerous previous studies to interact with the same N-terminal locus. Cellular Mg^2+^ may be altered in disease states; for example, it may be elevated in ischemic stress or decreased as in hyperparathyroidism. Mg^2+^-binding binding to cTnC may further alter the already skewed Ca^2+^-cTnC binding interaction which exists in diseases such as HCM or DCM, further affecting significant changes in cardiomyocyte E-C coupling.

## Author Contributions

Preliminary Experiments – KR

Experimental Design - KR, ERH, AMS, FvP, RJS, SL, GFT

Data Collection – KR, ERH

Data Analysis – KR, ERH, SL

Manuscript Preparation – KR, ERH, FvP, SL, GFT

Manuscript Review – KR, ERH, AMS, FvP, RJS, SL, GFT

## Experimental procedures

### Construct Preparations

Recombinant proteins were expressed and purified as described previously (96). In brief, the human cTnC gene (*TNNC1*) within the pET-21a(+) vector was ordered from Novagen and the Phusion site directed mutagenesis kit (Thermo) used to introduce a stop codon at residue 90, followed by single base pair changes to introduce all 6 mutations of interest (A8V, L29Q, A31S, L48Q, Q50R, and C84Y) on separate N-terminal constructs (cTnC_1-89_). Mutagenesis was carried out with preliminary steps using the DH5α *E. coli* strain to house the plasmids. Following the mutagenesis and confirmation by sequencing, the constructs were transformed into the BL21(DE3) expression strain and stored as glycerol stocks.

### Protein Expression

100 mL of Lysogeny Broth (LB) supplemented with 50 μg/mL of Ampicillin and a stab of the glycerol stock was grown overnight at 37 ° C for 16 – 20 hrs with shaking at 225-250 rpm. 1 L flasks of LB were induced with 1-5% of the over night culture and supplemented with the same concentration of antibiotic and grown under the same conditions for ~3 hrs (until OD600 was between 0.8-1.0). The culture was then supplemented with 1 mM Isopropyl β-D-1-thiogalactopyranoside (IPTG) and grown for a further 3 – 4 hrs. Cells were then harvested by centrifugation and resuspended in the Lysis Buffer (50 mM Tris-Cl and 100 mM NaCl at pH 8.0). The suspended pellet was stored at −80 ° C until purification.

### Protein Purification

The pellet was thawed and sonicated at ~80% amplitude in 30 second intervals for a total time of 3 – 4 mins with each intermittent period spent on ice. The cells were then spun 2 times, for 15 minutes each at 30,000 xg and the supernatant kept and the pellet discarded. The supernatant was filtered as needed and applied to a 15 mL fast-flow DEAE or Q Sepharose column (GE Healthcare), pre-equilibrated with Buffer A (50 mM Tris-Cl, 100 mM NaCl, and 1 mM Dithiothreitol (DTT) at pH 8.0). Buffer B (Buffer A + 0.55 M NaCl) was applied over a 180 mL protocol, where the concentration was ramped up from 0 to 100% to elute the proteins of interest. Fractions containing the N-terminal cTnC construct were identified by SDS PAGE and pooled. An Amicon centrifugal concentrator (Millipore) with a 3 KDa cut-off was used to concentrate the pooled samples to a volume of 3 – 5 mL. The pooled samples were then applied to a HiPrep 26/60 Sephacryl S-100 column (GE Healthcare) equilibrated with Buffer A. The fractions were again analyzed by SDS PAGE and those containing the protein of interest, free of contaminants were pooled, concentrated, and stored at −80 °C.

### ITC Experiments

The protein was dialyzed against 3 exchanges of 2 L for at least 6 hrs each with ITC Buffer 1 containing 50 mM HEPES, 150 mM KCl, 2 mM of EDTA, and 15 mM β-mercaptoethanol (BME) at pH 7.2. ITC Buffer 2 was identical to the first but did not contain EDTA. ITC Buffer 3 was identical to the second but contained 2 mM BME. The nanodrop was used to gain a preliminary measure of the protein concentration using an extinction coefficient of 1490 M^-1^*cm^-1^ and a molecular weight of 10.4 kDa. An initial ITC run, was used to determine the molar ratio (N). Given that the concentration of the titrant is known and the number of binding sites in N-cTnC is 1, the concentration of folded, functional protein can be determined and adjusted in subsequent runs to give an N of 1.0.

The protein was diluted in the final dialysis buffer to a final concentration of 100 μM. The titrating solutions were prepared from 1.00 M Ca^2+^ and Mg^2+^ stocks (Sigma) by serial dilution in the final dialysis buffer.

### ITC Protocol

For the apo-state experiments, 2 mM Ca^2+^ and 20 mM Mg^2+^ were titrated into 100 μM N-cTnC with the exception of the L48Q where 2 mM Mg^2+^ was used. For competition experiments, the Apo-state construct was pre-incubated with Mg^2+^ to a final concentration of 1 mM or 3 mM Mg^2+^ prior to titration with 2 mM Mg^2+^. Titrant into buffer blank experiments were carried out the gauge the impact of these experiments and indicate minimal heat change resulting from the interactions (**Figure S5**). For all experiments, 19 titrations, 60 seconds apart were performed with the first being 0.8 μL and each subsequent injection 2 μL. The cell contents were mixed at 750 rpm throughout the titration. All titrations were carried out at 25 ° C.

### Data Processing and Statistical Analysis

Data were imported and analyzed in Origin 8.0 software for Microcal ITC200 (Northampton, MA). After saturation, the final 2-3 data points were averaged, the heat was subtracted from all injections as a control for heat of dilution and non-specific interactions. Least-squares regression was used to fit each titration after the first (dummy) injection was removed with minimization of chi-square and visual evaluation used to determine the goodness-of-fit for a single binding site model. Following establishment of the protein concentration based on the obtained N value for each apo-state Ca^2+^ titration for each construct, the same dilution of protein was used for each other titration and the N-value fixed to 1.00 to facilitate data fitting. The various thermodynamic parameters were averaged and reported as a mean ± SEM. The difference between the means was compared using a one-way ANOVA, where all the titration conditions had p<0.001. This was followed by Tukey’s post-hoc test to determine where significant differences existed (p<0.05) (**Tables 1 and 2**).

### Thermodynamic Integration

The structure of the N-terminal domain of cardiac troponin C (N-cTnC) was obtained from PDB: 1AP4(97), this structure contained N-cTnC with a single Ca^2+^ ion bound. Since there was no model of Mg^2+^ bound N-cTnC in the protein databank, we made use of the Mg^2+^ substituted structure as outlined in our previous work (57). The model was then solvated using the tLeap module of AMBER 16(98) in a 12Å TIP3P water box and neutralized with Na^+^ ions; the forcefield used to describe the protein was ff14SB (99). In order to complete the thermodynamic cycle, a system containing the Mg^2+^ ion was prepared using the tLeap module, referencing the Δ*G_solvation_* optimized Mg^2+^ parameters from Li et. al. (100), and solvated in a 12Å TIP3P water box. Simulations were conducted under NPT conditions using the Berendsen barostat and periodic boundary conditions. The system was minimized for 2000 cycles and heated to 300 K using the Langevin thermostat over 500 ps prior to the 5 ns production with a time step of 2 fs. The SHAKE algorithm was employed to constrain all bonds involving hydrogen atoms, and the Particle Mesh Ewald method(101) was utilized to calculate electrostatic interactions of long distances with a cutoff of 10 Å.

The alchemical thermodynamic cycle used for ligand binding has been detailed previously by Leelananda and Lindert (102). In this implementation of TI, the method consisted of three steps for ligand (Mg^2+^) in protein: introduction of harmonic distance restraints, removal of electrostatic interactions, and removal of van der Waals forces. TI consisted of two steps for the ligand in water system: removal of electrostatic interactions and removal of van der Waals forces. The coupling parameter (λ) increased incrementally by 0.1 from 0.0 – 1.0 for each transitional step of the thermodynamic cycle. During each simulation dV/dλ values were collected every 2 ps resulting in 5000 data points per transitional step of λ for further analysis. The Multistate Bennett Acceptance Ratio (MBAR)(103) was used to calculate the relative free energies of the simulations across all values of λ. Free energy (Δ*G*) corrections were made due to the introduction of the distance restraints and to correct for the charge of the system as described previously(57). For each system, 5 independent runs were performed and the results averaged. The specific distance restraints for all protein structures are shown in **Table S2.**

Mutants (L48Q, Q50R, C84Y) were constructed using the protein mutagenesis tool in PyMOL(104) and the Mg^2+^ substituted representative model of N-cTnC serving as the base model. TI simulations were performed on the mutant structure as detailed above for the wild type.

## Supplementary Appendix

**Figure S1:**
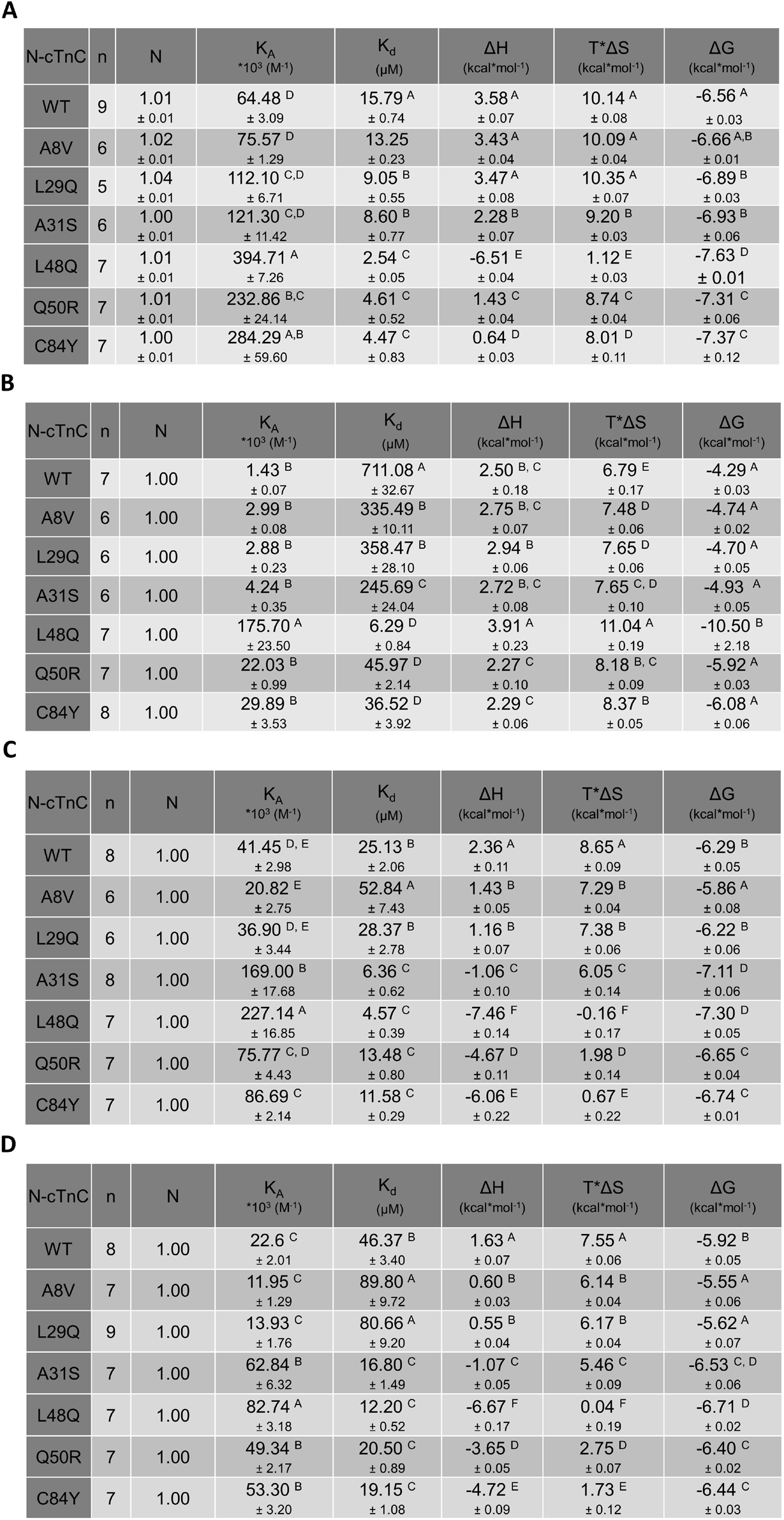
The thermodynamic properties of the binding interactions with WT N-cTnC and each of the mutants are listed. For each parameter and across the construct type, those differing in their mean p<0.05 from the WT are marked by *. Each parameter is displayed as mean ± SEM, with the exception of N which was fixed to 1.00 in the Mg^2+^ binding and preincubation experiments; A, the parameters associated with the titration of Ca^2+^ into apo-state N-cTnC; B, the parameters associated with the titration of Mg^2+^ into apo-state N-cTnC; C, the parameters associated with the titration of Ca^2+^ into 1 mM Mg^2+^ preincubated N-cTnC; D, the parameters associated with the titration of Ca^2+^ into 3 mM Mg^2+^ preincubated N-cTnC.

**Figure S1:**
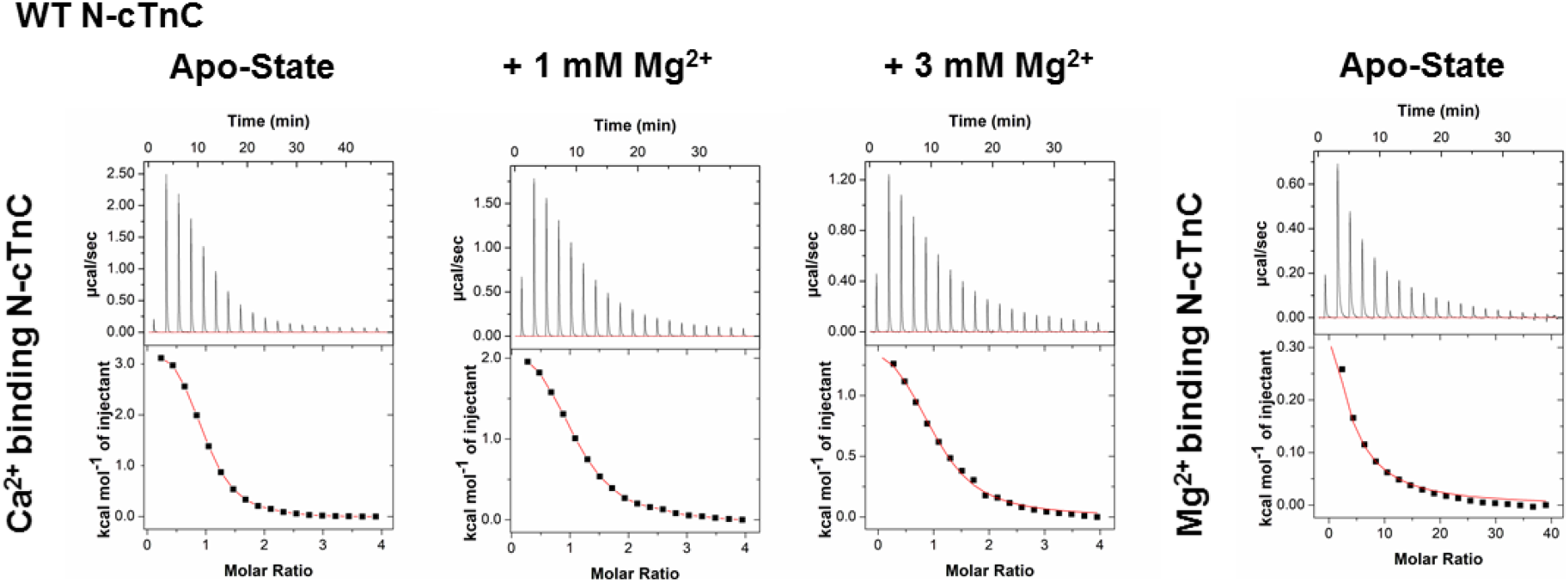
Representative isotherms for each titration condition in the WT N-cTnC construct. From the right, the titration of Ca^2+^ into apo-state N-cTnC is shown, the next two panels show the titration of Ca^2+^ into 1 mM and 3 mM Mg^2+^ incubated WT N-cTnC. The right-most panel shows titration of Mg^2+^ into apo-state protein. Each titration is similarly endothermic with the scales indicating differences in absolute value of change in enthalpy.

**Figure S2:**
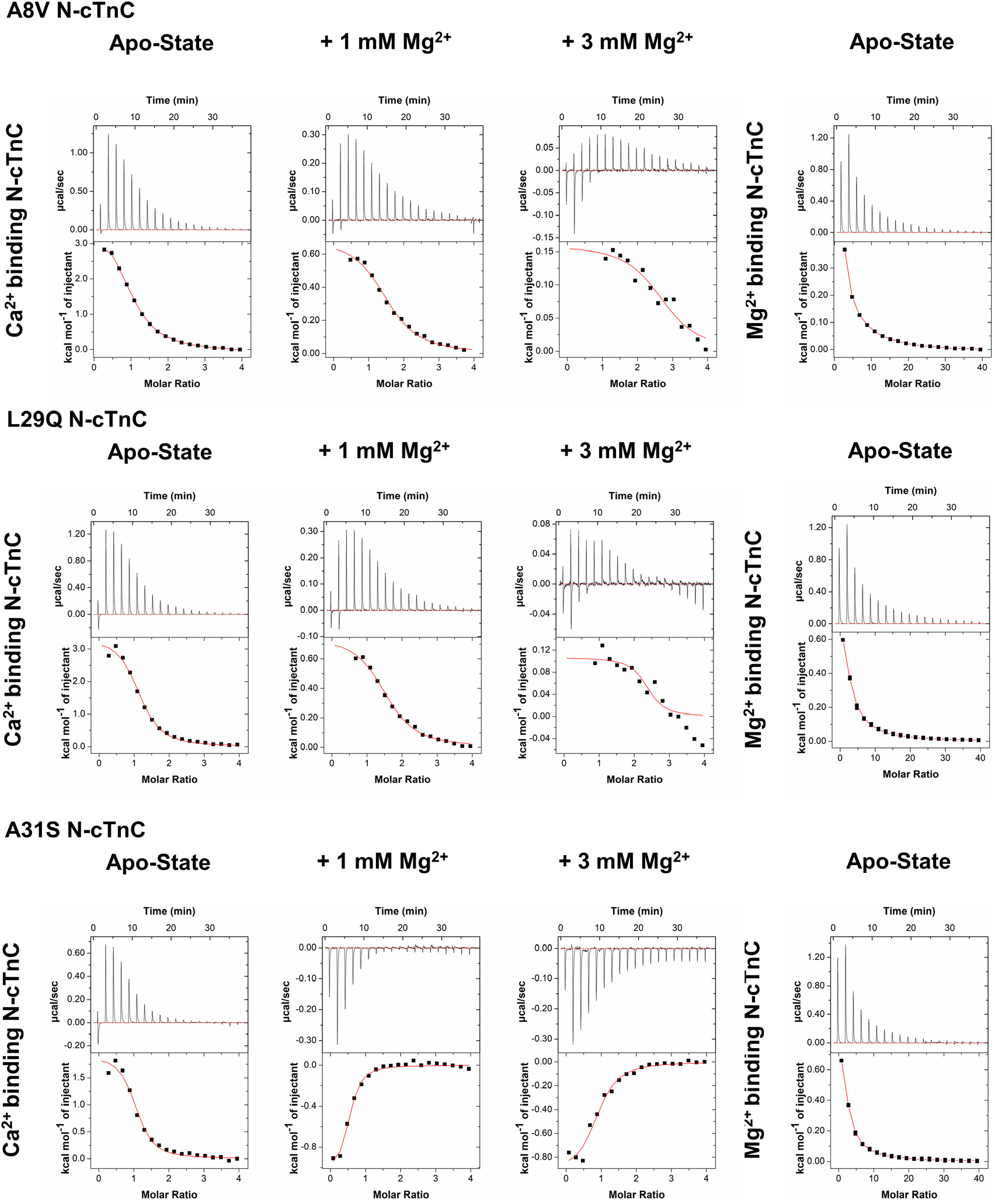
Representative isotherms for each titration condition into A8V, L29Q, and A31S N-cTnC. The 3 most N-terminal mutations are shown; A8V, L29Q, and A31S from top to bottom. On the left panel, titration of Ca^2+^ into apo-state protein is shown with the next panels show the titration of Ca^2+^ into 1 mM and 3 mM Mg^2+^ pre-incubated N-cTnC. The right most panel shows the titration of Mg^2+^ into apo-state N-cTnC. The majority of titrations are characterized by an endothermic interaction with the exception of A31S, where pre-incubation with Mg^2+^ resulted in an exothermic interaction with Ca^2+^.

**Figure S3:**
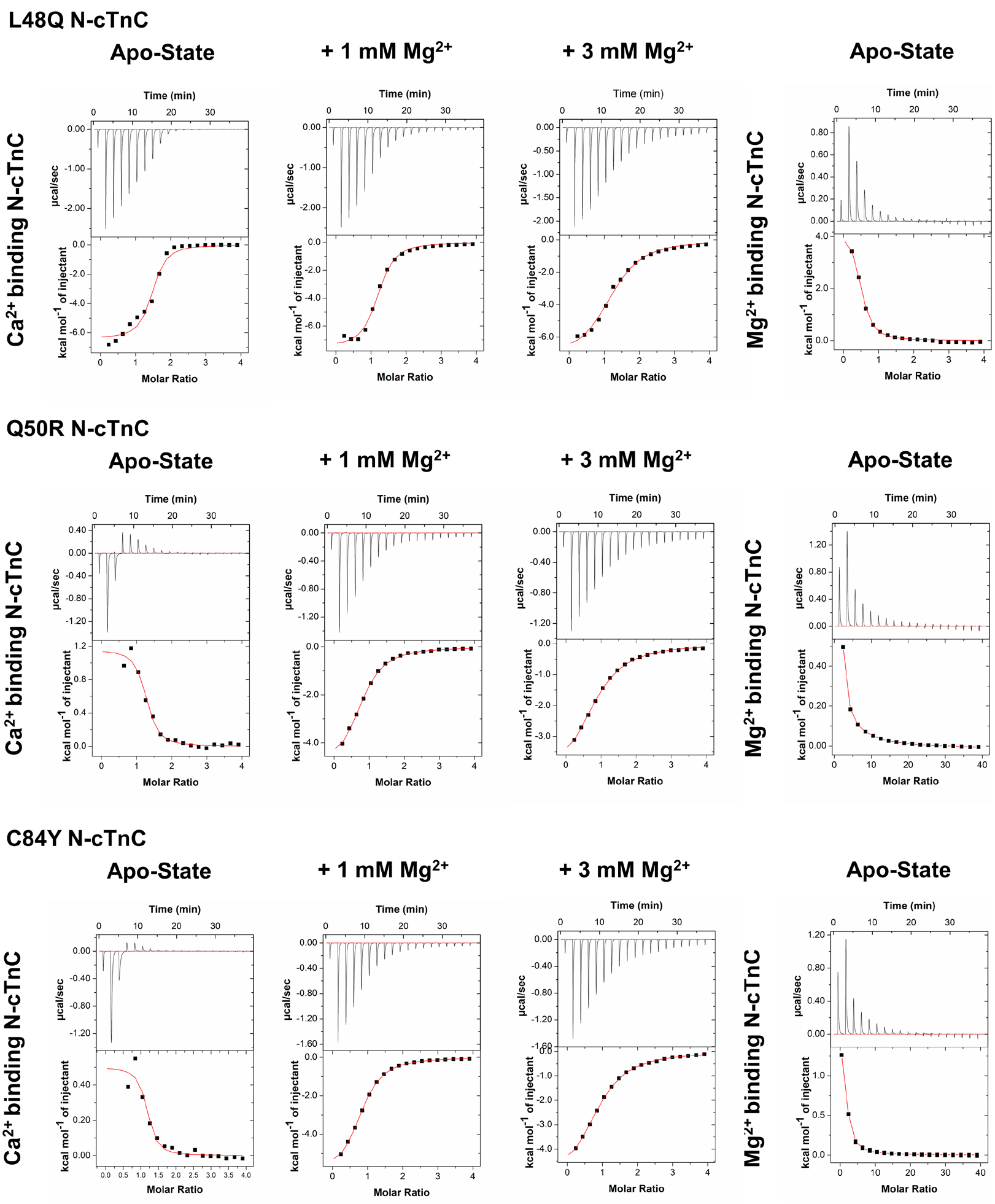
Representative isotherms for each titration condition for L48Q, Q50R, and, C84Y N-cTnC. In the 3 most N-terminal mutations are shown; L48Q, Q50R, and C84Y from top to bottom. On the left most panel, titration of Ca^2+^ into apo-state protein is shown with the next panels showing the titration of Ca^2+^ into 1 mM and 3 mM Mg^2+^ pre-incubated N-cTnC. The right most panel show the titration of Mg^2+^ into apo-state N-cTnC. These three mutants caused the greatest deviation in thermodynamic properties form the WT titration conditions. The Ca^2+^ into apo-protein titration is endothermic for Q50R and C84Y but exothermic for L48Q. The Mg^2+^ into apo-protein titration is endothermic for all 3 mutants. The pre-incubation condition with both 1 mM and 3 mM Mg^2+^ resulted in an exothermic interaction with Ca^2+^.

**Figure S5:**
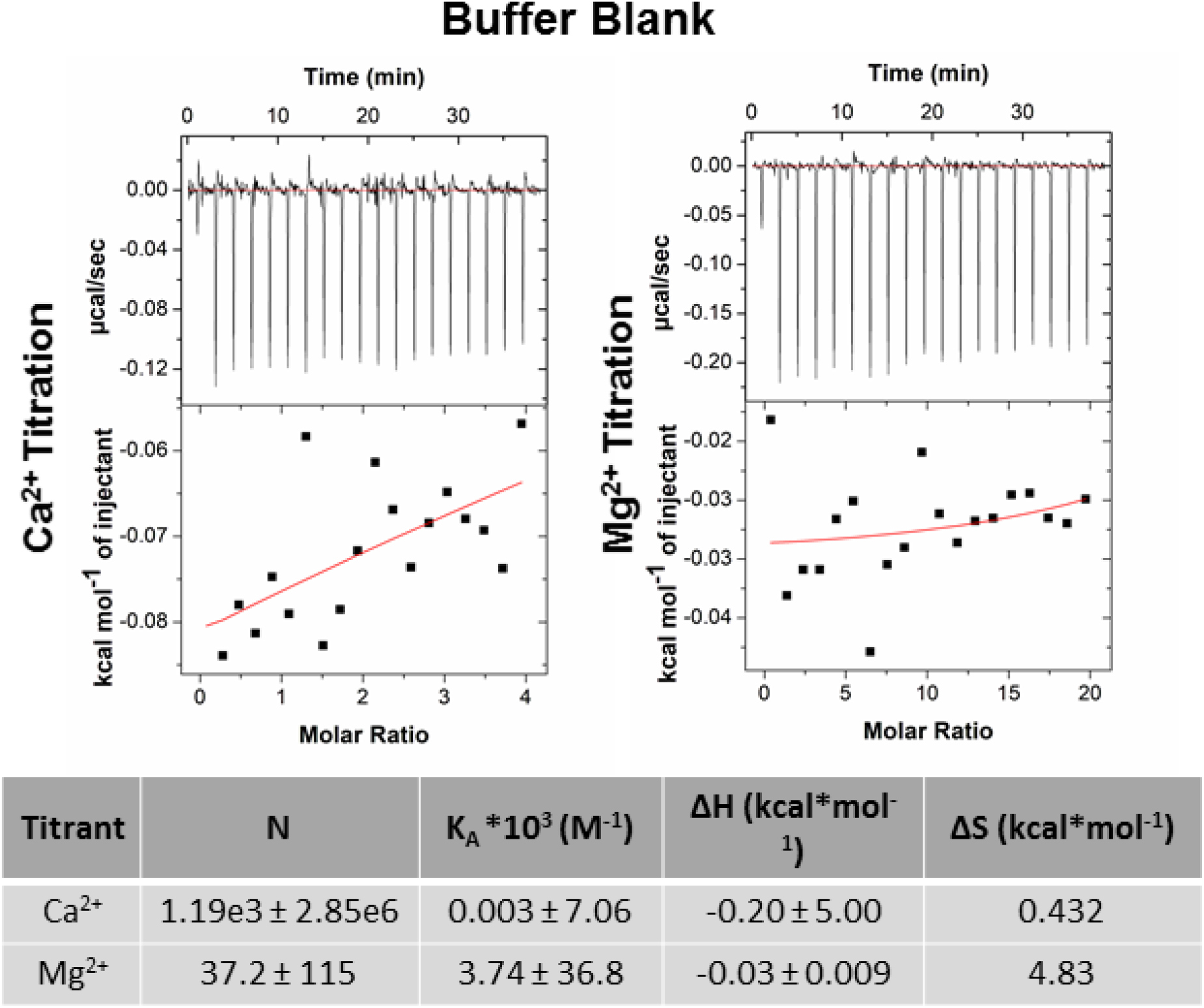
Representative isotherms are presented to quantify the heat signal from the interaction with each of Ca^2+^ and Mg^2+^ with the buffer. The scale may be used as measure of the heat change in the system and a proxy to compare with the titrant into N-cTnC conditions. The values measured when a single binding site model was used to fit each titration are presented in the table.

**Table S2.** Mg^2+^ restraint distances for thermodynamic integration

|  | WT N-cTnC |  | L48Q |  | Q50R |  | C84Y |  |
| --- | --- | --- | --- | --- | --- | --- | --- | --- |
| Restraint 1 | ASP67 CG<br>Atom 1029 | 2.3 Å | ASP67 CG<br>Atom 1027 | 2.3 Å | ASP67 CG<br>Atom 1036 | 2.3 Å | ASP67 CG<br>Atom 1029 | 2.3 Å |
| Restraint 2 | SER69 OG<br>Atom 1048 | 3.7 Å | SER69 OG<br>Atom 1046 | 3.7 Å | SER69 OG<br>Atom 1055 | 3.7 Å | SER69 OG<br>Atom 1048 | 3.7 Å |
| Restraint 3 | THR71 OG1<br>Atom 1067 | 4.5 Å | THR71 OG1<br>Atom 1065 | 4.5 Å | THR71 OG1<br>Atom 1074 | 4.5 Å | THR71 OG1<br>Atom 1067 | 4.5 Å |

## References

1. Potter, J. D., and Gergely, J. (1975) The calcium and magnesium binding sites on troponin and their role in the regulation of myofibrillar adenosine triphosphatase. Journal of Biological Chemistry 250, 4628–4633

2. Sturtevant, J. M. (1977) Heat capacity and entropy changes in processes involving proteins. Proc Natl Acad Sci U S A 74, 2236–2240

3. Tikunova, S. B., and Davis, J. P. (2004) Designing calcium-sensitizing mutations in the regulatory domain of cardiac troponin C. Journal of Biological Chemistry 279, 35341–35352

4. Bers, D. M. (2000) Calcium Fluxes Involved in Control of Cardiac Myocyte Contraction. Circulation Research 87, 275–281

5. Sia, S. K., Li, M. X., Spyracopoulos, L., Gagné, S. M., Liu, W., Putkey, J. A., and Sykes, B. D. (1997) Structure of cardiac muscle troponin C unexpectedly reveals a closed regulatory domain. Journal of Biological Chemistry 272, 18216–18221

6. Kirschenlohr, H. L., Grace, A. A., Vandenberg, J. I., Metcalfe, J. C., and Smith, G. A. (2000) Estimation of systolic and diastolic free intracellular Ca^2+^ by titration of Ca^2+^ buffering in the ferret heart. Biochem J 346 Pt 2, 385–391

7. Maron, B. J., Gardin, J. M., Flack, J. M., Gidding, S. S., Kurosaki, T. T., and Bild, D. E. (1995) Prevalence of hypertrophic cardiomyopathy in a general population of young adults Echocardiographic analysis of 4111 subjects in the CARDIA Study. Circulation 92, 785–789

8. Semsarian, C., Ingles, J., Maron, M. S., and Maron, B. J. (2015) New perspectives on the prevalence of hypertrophic cardiomyopathy. Journal of the American College of Cardiology 65, 1249–1254

9. Seidman, C. E., and Seidman, J. G. (2011) Identifying sarcomere gene mutations in hypertrophic cardiomyopathy: a personal history. Circ Res 108, 743–750

10. Ashrafian, H., and Watkins, H. (2007) Reviews of translational medicine and genomics in cardiovascular disease: new disease taxonomy and therapeutic implications: Cardiomyopathies: Therapeutics based on molecular phenotype. Journal of the American College of Cardiology 49, 1251–1264

11. Harada, K., and Morimoto, S. (2004) Inherited cardiomyopathies as a troponin disease. Jpn J Physiol 54, 307–318

12. Elliott, P., and McKenna, W. J. (2004) Hypertrophic cardiomyopathy. Lancet 363, 1881–1891

13. Goldspink, P. H., Warren, C. M., Kitajewski, J., Wolska, B. M., and Solaro, R. J. (2021) A perspective on personalized therapies in hypertrophic cardiomyopathy. Journal of Cardiovascular Pharmacology 77, 317–322

14. Maron, B. J., Shirani, J., Poliac, L. C., Mathenge, R., Roberts, W. C., and Mueller, F. O. (1996) Sudden death in young competitive athletes: clinical, demographic, and pathological profiles. JAMA 276, 199–204

15. Davis, J., Davis, L. C., Correll, R. N., Makarewich, C. A., Schwanekamp, J. A., Moussavi-Harami, F., Wang, D., York, A. J., Wu, H., and Houser, S. R. (2016) A tension-based model distinguishes hypertrophic versus dilated cardiomyopathy. Cell 165, 1147–1159

16. Katrukha, I. (2013) Human cardiac troponin complex. Structure and functions. Biochemistry (Moscow) 78, 1447–1465

17. Kalyva, A., Parthenakis, F. I., Marketou, M. E., Kontaraki, J. E., and Vardas, P. E. (2014) Biochemical characterisation of Troponin C mutations causing hypertrophic and dilated cardiomyopathies. Journal of muscle research and cell motility 35, 161–178

18. Cordina, N. M., Liew, C. K., Gell, D. A., Fajer, P. G., Mackay, J. P., and Brown, L. J. (2013) Effects of Calcium Binding and the Hypertrophic Cardiomyopathy A8V Mutation on the Dynamic Equilibrium between Closed and Open Conformations of the Regulatory N-Domain of Isolated Cardiac Troponin C. Biochemistry 52, 1950–1962

19. Hoffmann, B., Schmidt-Traub, H., Perrot, A., Osterziel, K. J., and Gessner, R. (2001) First mutation in cardiac troponin C, L29Q, in a patient with hypertrophic cardiomyopathy. Hum Mutat 17, 524

20. Parvatiyar, M. S., Landstrom, A. P., Figueiredo-Freitas, C., Potter, J. D., Ackerman, M. J., and Pinto, J. R. (2012) A mutation in TNNC1-encoded cardiac troponin C, TNNC1-A31S, predisposes to hypertrophic cardiomyopathy and ventricular fibrillation. Journal of Biological Chemistry 287, 31845–31855

21. Landstrom, A. P., Parvatiyar, M. S., Pinto, J. R., Marquardt, M. L., Bos, J. M., Tester, D. J., Ommen, S. R., Potter, J. D., and Ackerman, M. J. (2008) Molecular and functional characterization of novel hypertrophic cardiomyopathy susceptibility mutations in TNNC1-encoded troponin C. Journal of molecular and cellular cardiology 45, 281–288

22. van Spaendonck-Zwarts, K. Y., van Tintelen, J. P., van Veldhuisen, D. J., van der Werf, R., Jongbloed, J. D., Paulus, W. J., Dooijes, D., and van den Berg, M. P. (2010) Peripartum cardiomyopathy as a part of familial dilated cardiomyopathy. Circulation 121, 2169–2175

23. Stevens, C. M., Rayani, K., Singh, G., Lotfalisalmasi, B., Tieleman, D. P., and Tibbits, G. F. (2017) Changes in the dynamics of the cardiac troponin C molecule explain the effects of Ca2+-sensitizing mutations. Journal of Biological Chemistry 292, 11915–11926

24. Gillis, T. E., Blumenschein, T. M., Sykes, B. D., and Tibbits, G. F. (2003) Effect of temperature and the F27W mutation on the Ca^2+^ activated structural transition of trout cardiac troponin C. Biochemistry 42, 6418–6426

25. Gillis, T. E., Liang, B., Chung, F., and Tibbits, G. F. (2005) Increasing mammalian cardiomyocyte contractility with residues identified in trout troponin C. Physiological genomics 22, 1–7

26. Gillis, T. E., Moyes, C. D., and Tibbits, G. F. (2003) Sequence mutations in teleost cardiac troponin C that are permissive of high Ca^2+^ affinity of site II. American Journal of Physiology-Cell Physiology 284, C1176–C1184

27. Davis, J. P., Rall, J. A., Reiser, P. J., Smillie, L. B., and Tikunova, S. B. (2002) Engineering competitive magnesium binding into the first EF-hand of skeletal troponin C. The Journal of biological chemistry 277, 49716–49726

28. Li, A. Y., Stevens, C. M., Liang, B., Rayani, K., Little, S., Davis, J., and Tibbits, G. F. (2013) Familial hypertrophic cardiomyopathy related cardiac troponin C L29Q mutation alters length-dependent activation and functional effects of phosphomimetic troponin I*.

29. Yamada, K. (2003) Calcium binding to troponin C as a primary step of the regulation of contraction. A microcalorimetric approach. Adv Exp Med Biol 538, 203–212; discussion 213

30. Wilcox, D. E. (2008) Isothermal titration calorimetry of metal ions binding to proteins: An overview of recent studies. Inorganica Chimica Acta 361, 857–867

31. Grossoehme, N. E., Spuches, A. M., and Wilcox, D. E. (2010) Application of isothermal titration calorimetry in bioinorganic chemistry. J Biol Inorg Chem 15, 1183–1191

32. Sacco, C., Skowronsky, R. A., Gade, S., Kenney, J. M., and Spuches, A. M. (2012) Calorimetric investigation of copper(II) binding to Abeta peptides: thermodynamics of coordination plasticity. J Biol Inorg Chem 17, 531–541

33. Skowronsky, R. A., Schroeter, M., Baxley, T., Li, Y., Chalovich, J. M., and Spuches, A. M. (2013) Thermodynamics and molecular dynamics simulations of calcium binding to the regulatory site of human cardiac troponin C: evidence for communication with the structural calcium binding sites. JBIC Journal of Biological Inorganic Chemistry 18, 49–58

34. Freire, E., Mayorga, O. L., and Straume, M. (1990) Isothermal titration calorimetry. Analytical chemistry 62, 950A–959A

35. Dai, L. J., Friedman, P. A., and Quamme, G. A. (1997) Phosphate depletion diminishes Mg2+ uptake in mouse distal convoluted tubule cells. Kidney Int 51, 1710–1718

36. Romani, A., and Scarpa, A. (1992) Regulation of cell magnesium. Arch Biochem Biophys 298, 1–12

37. Maguire, M. E. (2006) Magnesium transporters: properties, regulation and structure. Front Biosci 11, 3149–3163

38. Krause, S. M., and Rozanski, D. (1991) Effects of an increase in intracellular free [Mg2+] after myocardial stunning on sarcoplasmic reticulum Ca2+ transport. Circulation 84, 1378–1383

39. Kirkels, J., Van Echteld, C., and Ruigrok, T. (1989) Intracellular magnesium during myocardial ischemia and reperfusion: possible consequences for postischemic recovery. Journal of molecular and cellular cardiology 21, 1209–1218

40. Potter, J. D., Robertson, S. P., and Johnson, J. D. (1981) Magnesium and the regulation of muscle contraction. Fed Proc 40, 2653–2656

41. Ogawa, Y. (1985) Calcium binding to troponin C and troponin: effects of Mg2+, ionic strength and pH. The Journal of Biochemistry 97, 1011–1023

42. Zot, A. S., and Potter, J. D. (1987) Structural aspects of troponin-tropomyosin regulation of skeletal muscle contraction. Annual review of biophysics and biophysical chemistry 16, 535–559

43. Morimoto, S. (1991) Effect of myosin cross-bridge interaction with actin on the Ca2+-binding properties of troponin C in fast skeletal myofibrils. The Journal of Biochemistry 109, 120–126

44. Francois, J. M., Gerday, C., Prendergast, F. G., and Potter, J. D. (1993) Determination of the Ca^2+^ and Mg^2+^ affinity constants of troponin C from eel skeletal muscle and positioning of the single tryptophan in the primary structure. J Muscle Res Cell Motil 14, 585–593

45. She, M., Dong, W. J., Umeda, P. K., and Cheung, H. C. (1998) Tryptophan mutants of troponin C from skeletal muscle: an optical probe of the regulatory domain. European journal of biochemistry 252, 600–607

46. Allen, T. S., Yates, L. D., and Gordon, A. M. (1992) Ca^2+^-dependence of structural changes in troponin-C in demembranated fibers of rabbit psoas muscle. Biophys J 61, 399–409

47. Godt, R. E., and Morgan, J. L. (1984) Contractile responses to MgATP and pH in a thick filament regulated muscle: studies with skinned scallop fibers. Adv Exp Med Biol 170, 569–572

48. Godt, R. E. (1974) Calcium-activated tension of skinned muscle fibers of the frog. Dependence on magnesium adenosine triphosphate concentration. J Gen Physiol 63, 722–739

49. Best, P. M., Donaldson, S. K., and Kerrick, W. G. (1977) Tension in mechanically disrupted mammalian cardiac cells: effects of magnesium adenosine triphosphate. J Physiol 265, 1–17

50. Willott, R. H., Gomes, A. V., Chang, A. N., Parvatiyar, M. S., Pinto, J. R., and Potter, J. D. (2010) Mutations in Troponin that cause HCM, DCM AND RCM: what can we learn about thin filament function? Journal of molecular and cellular cardiology 48, 882–892

51. Liang, B., Chung, F., Qu, Y., Pavlov, D., Gillis, T. E., Tikunova, S. B., Davis, J. P., and Tibbits, G. F. (2008) Familial hypertrophic cardiomyopathy-related cardiac troponin C mutation L29Q affects Ca^2+^ binding and myofilament contractility. Physiological genomics 33, 257–266

52. Tikunova, S. B., Liu, B., Swindle, N., Little, S. C., Gomes, A. V., Swartz, D. R., and Davis, J. P. (2010) Effect of calcium-sensitizing mutations on calcium binding and exchange with troponin C in increasingly complex biochemical systems. Biochemistry 49, 1975–1984

53. Gomes, A. V., and Potter, J. D. (2004) Molecular and cellular aspects of troponin cardiomyopathies. Ann N Y Acad Sci 1015, 214–224

54. Ohki, S., Ikura, M., and Zhang, M. (1997) Identification of Mg2+-binding sites and the role of Mg2+ on target recognition by calmodulin. Biochemistry 36, 4309–4316

55. Malmendal, A., Evenas, J., Thulin, E., Gippert, G. P., Drakenberg, T., and Forsen, S. (1998) When size is important. Accommodation of magnesium in a calcium binding regulatory domain. The Journal of biological chemistry 273, 28994–29001

56. Andersson, M., Malmendal, A., Linse, S., Ivarsson, I., Forsen, S., and Svensson, L. A. (1997) Structural basis for the negative allostery between Ca(2+)- and Mg(2+)-binding in the intracellular Ca(2+)-receptor calbindin D9k. Protein Sci 6, 1139–1147

57. Rayani, K., Seffernick, J., Li, A. Y., Davis, J. P., Spuches, A. M., Van Petegem, F., Solaro, R. J., Lindert, S., and Tibbits, G. F. (2021) Binding of calcium and magnesium to human cardiac Troponin C. The Journal of biological chemistry, 100350

58. Gillis, T. E., Marshall, C. R., and Tibbits, G. F. (2007) Functional and evolutionary relationships of troponin C. Physiol Genomics 32, 16–27

59. Yamada, Y., Namba, K., and Fujii, T. (2020) Cardiac muscle thin filament structures reveal calcium regulatory mechanism. Nature communications 11, 1–9

60. Greenberg, M. J., and Tardiff, J. C. (2021) Complexity in genetic cardiomyopathies and new approaches for mechanism-based precision medicine. Journal of General Physiology 153

61. Tobacman, L. S., and Cammarato, A. (2021) Cardiomyopathic troponin mutations predominantly occur at its interface with actin and tropomyosin. Journal of General Physiology 153

62. Dong, W.-J., Wang, C.-K., Gordon, A. M., and Cheung, H. C. (1997) Disparate Fluorescence Properties of 2-[4’-(Iodoacetamido)anilino]-Naphthalene-6-Sulfonic Acid Attached to Cys-84 and Cys-35 of Troponin C in Cardiac Muscle Troponin. Biophysical Journal 72, 850–857

63. Holroyde, M., Robertson, S., Johnson, J., Solaro, R., and Potter, J. (1980) The calcium and magnesium binding sites on cardiac troponin and their role in the regulation of myofibrillar adenosine triphosphatase. Journal of Biological Chemistry 255, 11688–11693

64. Ebashi, S., and Ogawa, Y. (1988) Ca^2+^ in contractile processes. Biophys Chem 29, 137–143

65. Fabiato, A., and Fabiato, F. (1975) Effects of magnesium on contractile activation of skinned cardiac cells. J Physiol 249, 497–517

66. Donaldson, S. K., and Kerrick, W. G. (1975) Characterization of the effects of Mg^2+^ on Ca^2+^- and Sr^2+^-activated tension generation of skinned skeletal muscle fibers. J Gen Physiol 66, 427–444

67. Kerrick, W. G., Zot, H. G., Hoar, P. E., and Potter, J. D. (1985) Evidence that the Sr2+ activation properties of cardiac troponin C are altered when substituted into skinned skeletal muscle fibers. The Journal of biological chemistry 260, 15687–15693

68. Solaro, R. J., and Shiner, J. S. (1976) Modulation of Ca^2+^ control of dog and rabbit cardiac myofibrils by Mg^2+^. Comparison with rabbit skeletal myofibrils. Circ Res 39, 8–14

69. Ashley, C. C., and Moisescu, D. G. (1977) Effect of changing the composition of the bathing solutions upon the isometric tension-pCa relationship in bundles of crustacean myofibrils. J Physiol 270, 627–652

70. Donaldson, S. K., Best, P. M., and Kerrick, G. L. (1978) Characterization of the effects of Mg^2+^ on Ca^2+^- and Sr^2+^-activated tension generation of skinned rat cardiac fibers. J Gen Physiol 71, 645–655

71. Tanaka, H., Takahashi, H., and Ojima, T. (2013) Ca^2+^-binding properties and regulatory roles of lobster troponin C sites II and IV. FEBS Lett 587, 2612–2616

72. Pinto, J. R., Parvatiyar, M. S., Jones, M. A., Liang, J., Ackerman, M. J., and Potter, J. D. (2009) A functional and structural study of troponin C mutations related to hypertrophic cardiomyopathy. The Journal of biological chemistry 284, 19090–19100

73. Ramos, C. H. (1999) Mapping subdomains in the C-terminal region of troponin I involved in its binding to troponin C and to thin filament. The Journal of biological chemistry 274, 18189–18195

74. Gifford, Jessica L., Walsh, Michael P., and Vogel, Hans J. (2007) Structures and metal-ion-binding properties of the Ca^2+^-binding helix–loop–helix EF-hand motifs. Biochemical Journal 405, 199–221

75. Bowman, J. D., and Lindert, S. (2018) Molecular Dynamics and Umbrella Sampling Simulations Elucidate Differences in Troponin C Isoform and Mutant Hydrophobic Patch Exposure. 122, 7874–7883

76. Zot, H. G., Hasbun, J. E., Michell, C. A., Landim-Vieira, M., and Pinto, J. R. (2016) Enhanced troponin I binding explains the functional changes produced by the hypertrophic cardiomyopathy mutation A8V of cardiac troponin C. Arch Biochem Biophys 601, 97–104

77. Dweck, D., Hus, N., and Potter, J. D. (2008) Challenging current paradigms related to cardiomyopathies Are changes in the Ca2+ sensitivity of myofilaments containing cardiac troponin C mutations (G159D and L29Q) good predictors of the phenotypic outcomes? Journal of Biological Chemistry 283, 33119–33128

78. Pearlstone, J. R., Borgford, T., Chandra, M., Oikawa, K., Kay, C. M., Herzberg, O., Moult, J., Herklotz, A., Reinach, F. C., and Smillie, L. B. (1992) Construction and characterization of a spectral probe mutant of troponin C: application to analyses of mutants with increased calcium affinity. Biochemistry 31, 6545–6553

79. da Silva, A. C., de Araujo, A. H., Herzberg, O., Moult, J., Sorenson, M., and Reinach, F. C. (1993) Troponin-C mutants with increased calcium affinity. Eur J Biochem 213, 599–604

80. Potluri, P. R., Cordina, N. M., Kachooei, E., and Brown, L. J. (2019) Characterization of the L29Q Hypertrophic Cardiomyopathy Mutation in Cardiac Troponin C by Paramagnetic Relaxation Enhancement Nuclear Magnetic Resonance. 58, 908–917

81. Johnson, J. D., Collins, J. H., and Potter, J. D. (1978) Dansylaziridine-labeled troponin C. A fluorescent probe of Ca2+ binding to the Ca2+-specific regulatory sites. The Journal of biological chemistry 253, 6451–6458

82. Ward, W. H., and Holdgate, G. A. (2001) 7 Isothermal Titration Calorimetry in Drug Discovery. in Progress in medicinal chemistry, Elsevier. pp 309–376

83. Wang, D., Robertson, I. M., Li, M. X., McCully, M. E., Crane, M. L., Luo, Z., Tu, A.-Y., Daggett, V., Sykes, B. D., and Regnier, M. (2012) Structural and functional consequences of the cardiac troponin C L48Q Ca2+-sensitizing mutation. Biochemistry 51, 4473–4487

84. Shettigar, V., Zhang, B., Little, S. C., Salhi, H. E., Hansen, B. J., Li, N., Zhang, J., Roof, S. R., Ho, H. T., Brunello, L., Lerch, J. K., Weisleder, N., Fedorov, V. V., Accornero, F., Rafael-Fortney, J. A., Gyorke, S., Janssen, P. M., Biesiadecki, B. J., Ziolo, M. T., and Davis, J. P. (2016) Rationally engineered Troponin C modulates in vivo cardiac function and performance in health and disease. Nat Commun 7, 10794

85. Cummins, P. L., and Gready, J. E. (1994) Thermodynamic integration calculations on the relative free energies of complex ions in aqueous solution: Application to ligands of dihydrofolate reductase. Journal of Computational Chemistry 15, 704–718

86. Panteva, M. T., Giambaşu, G. M., and York, D. M. (2015) Comparison of structural, thermodynamic, kinetic and mass transport properties of Mg(2+) ion models commonly used in biomolecular simulations. J Comput Chem 36, 970–982

87. Reid, R. E., and Procyshyn, R. M. (1995) Engineering magnesium selectivity in the helix-loop-helix calcium-binding motif. Arch Biochem Biophys 323, 115–119

88. Tikunova, S. B., Black, D. J., Johnson, J. D., and Davis, J. P. (2001) Modifying Mg^2+^ binding and exchange with the N-terminal of calmodulin. Biochemistry 40, 3348–3353

89. Dai, L. J., Friedman, P. A., and Quamme, G. A. (1997) Cellular mechanisms of chlorothiazide and cellular potassium depletion on Mg2+ uptake in mouse distal convoluted tubule cells. Kidney Int 51, 1008–1017

90. Bers, D. M. (2002) Cardiac excitation–contraction coupling. Nature 415, 198–205

91. Touyz, R. M. (2004) Reactive oxygen species, vascular oxidative stress, and redox signaling in hypertension: what is the clinical significance? Hypertension 44, 248–252

92. Weglicki, W., Quamme, G., Tucker, K., Haigney, M., and Resnick, L. (2005) Potassium, magnesium, and electrolyte imbalance and complications in disease management. Clin Exp Hypertens 27, 95–112

93. Mazur, A., Maier, J. A., Rock, E., Gueux, E., Nowacki, W., and Rayssiguier, Y. (2007) Magnesium and the inflammatory response: potential physiopathological implications. Arch Biochem Biophys 458, 48–56

94. Kolte, D., Vijayaraghavan, K., Khera, S., Sica, D. A., and Frishman, W. H. (2014) Role of magnesium in cardiovascular diseases. Cardiology in review 22, 182–192

95. Murphy, E., Steenbergen, C., Levy, L. A., Raju, B., and London, R. E. (1989) Cytosolic free magnesium levels in ischemic rat heart. The Journal of biological chemistry 264, 5622–5627

96. Stevens, C. M., Rayani, K., Genge, C. E., Singh, G., Liang, B., Roller, J. M., Li, C., Li, A. Y., Tieleman, D. P., and van Petegem, F. (2016) Characterization of Zebrafish Cardiac and Slow Skeletal Troponin C Paralogs by MD Simulation and ITC. Biophysical Journal 111, 38–49

97. Spyracopoulos, L., Li, M. X., Sia, S. K., Gagné, S. M., Chandra, M., Solaro, R. J., and Sykes, B. D. (1997) Calcium-Induced Structural Transition in the Regulatory Domain of Human Cardiac Troponin C. Biochemistry 36, 12138–12146

98. Case, D. A., Betz, R. M., Cerutti, D. S., Cheatham, I., T.E., Darden, T. A., Duke, R. E., Giese, T. J., Gohlke, H., Goetz, A. W., Homeyer, N., Izadi, S., Janowski, P., Kaus, J., Kovalenko, A., Lee, T. S., LeGrand, S., Li, P., Lin, C., Luchko, T., Luo, R., Madej, B., Mermelstein, D., Merz, K. M., Monard, G., Nguyen, H., Nguyen, H. T., Omelyan, I., Onufriev, A., Roe, D. R., Roitberg, A., Sagui, C., Simmerling, C. L., Botello-Smith, W. M., Swails, J., Walker, R. C., Wang, J., Wolf, R. M., Wu, X., Xiao, L., and Kollman, P. A. (2016) AMBER 2016. University of California, San Francisco

99. Maier, J. A., Martinez, C., Kasavajhala, K., Wickstrom, L., Hauser, K. E., and Simmerling, C. (2015) ff14SB: Improving the Accuracy of Protein Side Chain and Backbone Parameters from ff99SB. Journal of Chemical Theory and Computation 11, 3696–3713

100. Li, P., Roberts, B. P., Chakravorty, D. K., and Merz, K. M. (2013) Rational Design of Particle Mesh Ewald Compatible Lennard-Jones Parameters for +2 Metal Cations in Explicit Solvent. J Chem Theory Comput 9, 2733–2748

101. Essmann, U., Perera, L., Berkowitz, M. L., Darden, T., Lee, H., and Pedersen, L. G. (1995) A smooth particle mesh Ewald method. The Journal of Chemical Physics 103, 8577–8593

102. Leelananda, S. P., and Lindert, S. (2016) Computational methods in drug discovery. Beilstein J Org Chem 12, 2694–2718

103. Shirts, M. R., and Chodera, J. D. (2008) Statistically optimal analysis of samples from multiple equilibrium states. J Chem Phys 129, 124105

104. The PyMOL Molecular Graphics System. Version 2.0 Ed., Schrödinger, LLC

